# Skeletal geometry and niche transitions restore organ size and shape during zebrafish fin regeneration

**DOI:** 10.1101/606970

**Authors:** Scott Stewart, Gabriel A. Yette, Heather K. Le Bleu, Astra L. Henner, Joshua A. Braunstein, Jad W. Chehab, Michael J. Harms, Kryn Stankunas

## Abstract

Regenerating fish fins return to their original size and shape regardless of the nature or extent of injury. Prevailing models for this longstanding mystery of appendage regeneration speculate fin cells maintain uncharacterized positional identities that instruct outgrowth after injury. Using zebrafish, we find differential Wnt production correlates with the extent of regeneration across the caudal fin. We identify Dachshund transcription factors as markers of distal blastema cells that produce Wnt and thereby promote a pro-progenitor and -proliferation environment. We show these Dach-expressing “niche cells” derive from mesenchyme populating cylindrical and progressively tapered fin rays. The niche pool, and consequently Wnt, steadily dissipates as regeneration proceeds; once exhausted, ray and fin growth stops. Supported by mathematical modeling, we show *longfin^t2^* zebrafish regenerate exceptionally long fins due to a perdurant niche, representing a “broken countdown timer”. We propose regenerated fin size is dictated by the amount of niche formed upon damage, which simply depends on the availability of intra-ray mesenchyme defined by skeletal girth at the injury site. Likewise, the fin reestablishes a tapered ray skeleton because progenitor osteoblast output reflects diminishing niche size. This “transpositional scaling” model contends mesenchyme-niche state transitions and positional information provided by self-restoring skeletal geometry rather than cell memories determine a regenerated fin’s size and shape.

## MAIN TEXT

Regenerating organs restore their original size and shape after injury. Vertebrate appendage regeneration, including that of teleost fish fins, provides a striking example of this phenomenon. Major fin amputations, tiny resections, and cuts of diverse geometry all produce the same outcome – a restored fin matching the original’s form and in scale with the animal’s body. Spallanzani, Broussonet, and T. H. Morgan pioneered studies of this longstanding mystery of regeneration in the 18^th^ and 19^th^ centuries (Broussonet, 1786; Morgan, 1900). For example, Morgan used oblique caudal fin resections to show that regeneration rates initially correlate with the amount of tissue lost and then progressively slow, ultimately stopping growth when the original size is regained (Morgan, 1900).

Prevailing models, now largely from zebrafish studies, posit that fin tissue adjacent to damage sites interpret Cartesian coordinate-like positional information to trigger the appropriate rate and extent of re-growth (Nachtrab et al., 2013; Rabinowitz et al., 2017; Rolland-Lagan et al., 2012; Tornini et al., 2016; Wolpert, 2016). This concept supposes an extensive array of positional identities, some epigenetic mechanism for cells to store such identities, and, perhaps most puzzling, for the positional information to be restored during the regeneration process. Ultimately, however, the rate and extent of outgrowth is dictated by production of growth factors, including FGF (Lee et al., 2005; Poss et al., 2000; Shibata et al., 2016) and Wnt (Stewart et al., 2014; Stoick-Cooper et al., 2007; Wehner et al., 2014), with the net effect of promoting cellular proliferation. Therefore, understanding growth factor production dynamics provides a logical entry to uncover fin size and shape restoration mechanisms.

Progressive fin regenerative outgrowth depends on spatially segregated growth factor production within an organized blastema. The blastema comprises heterogeneous cell types and forms soon after amputation by de-differentiation of mature cells and their migration into a cavity defined by the enveloping wound epidermis (Wehner and Weidinger, 2015). The blastema then arranges by both cell type and cell state, with distal progenitor and proximal differentiating zones. De-differentiated progenitor osteoblasts (pObs) migrate to blastema peripheries and hierarchically arrange with most progenitor state cells distally enriched (Stewart et al., 2014). Differentiating osteoblasts derived from the pOb pool locate more proximally and continue maturing to progressively extend re-forming bony rays. The central core of the blastema is largely comprised of “activated” mesenchymal cells that spill out from injury-exposed intra-ray space (Tornini et al., 2016; Tornini et al., 2017). These mesenchymal cells are the likely source of a distal, specialized “orchestrating center” (Wehner et al., 2014) that generates a pro-growth environment, including by producing mitogenic FGF (Lee et al., 2005; Poss et al., 2000; Shibata et al., 2016) and pro-progenitor Wnt (Lee et al., 2009; Stewart et al., 2014; Stoick-Cooper et al., 2007; Wehner et al., 2014). An even further distal, small pool of distinct blastema cells may serve as a largely quiescent reservoir (Nechiporuk and Keating, 2002). Distal blastema progenitor-supporting activity is balanced by proximal pro-differentiation signals, including BMP and Wnt-opposing Dkk1 produced by pObs that transitioned from a progenitor to differentiating state (Stewart et al., 2014).

Wnt signals indirectly promote cell proliferation (Wehner et al., 2014), in part by maintaining pObs (Stewart et al., 2014), to drive outgrowth until the fin is fully restored. Wnts such as *wnt5a*, *wnt5b*, and *wnt10a* are produced in the regenerating fin (Lee et al., 2009; Stewart et al., 2014; Stoick-Cooper et al., 2007; Wehner et al., 2014) including by distal blastema cells. We hypothesized differential Wnt production determines the rate and ultimately extent of regeneration. Revisiting Morgan’s experiments (Morgan, 1900), we obliquely amputated adult zebrafish fins from the distal tip of the dorsal side to a proximal location ventrally (see Fig. S1 for fin amputation planes and anatomical definitions). We then variably inhibited Wnt secretion by graded dosing of low concentrations of Wnt-C59 (Proffitt et al., 2013; Stewart et al., 2014). Partial perturbation of Wnt signaling prevented dorsal but not ventral tissue regeneration (Fig. 1A-E). Similarly, central tissue was more sensitive than peripheral tissue to Wnt inhibition following perpendicular resections (Fig. S2). We infer differential Wnt production underlies reacquisition of the stereotypical shape of a zebrafish fin. Concordantly, levels of *wnt5a*, a representative distal blastema-expressed Wnt (Lee et al., 2009; Stewart et al., 2014; Stoick-Cooper et al., 2007; Wehner et al., 2014), correlated with the demand for regenerative growth along the proximal-distal axis four days after oblique fin amputations (Fig. 1F, G).

**Figure 1.**
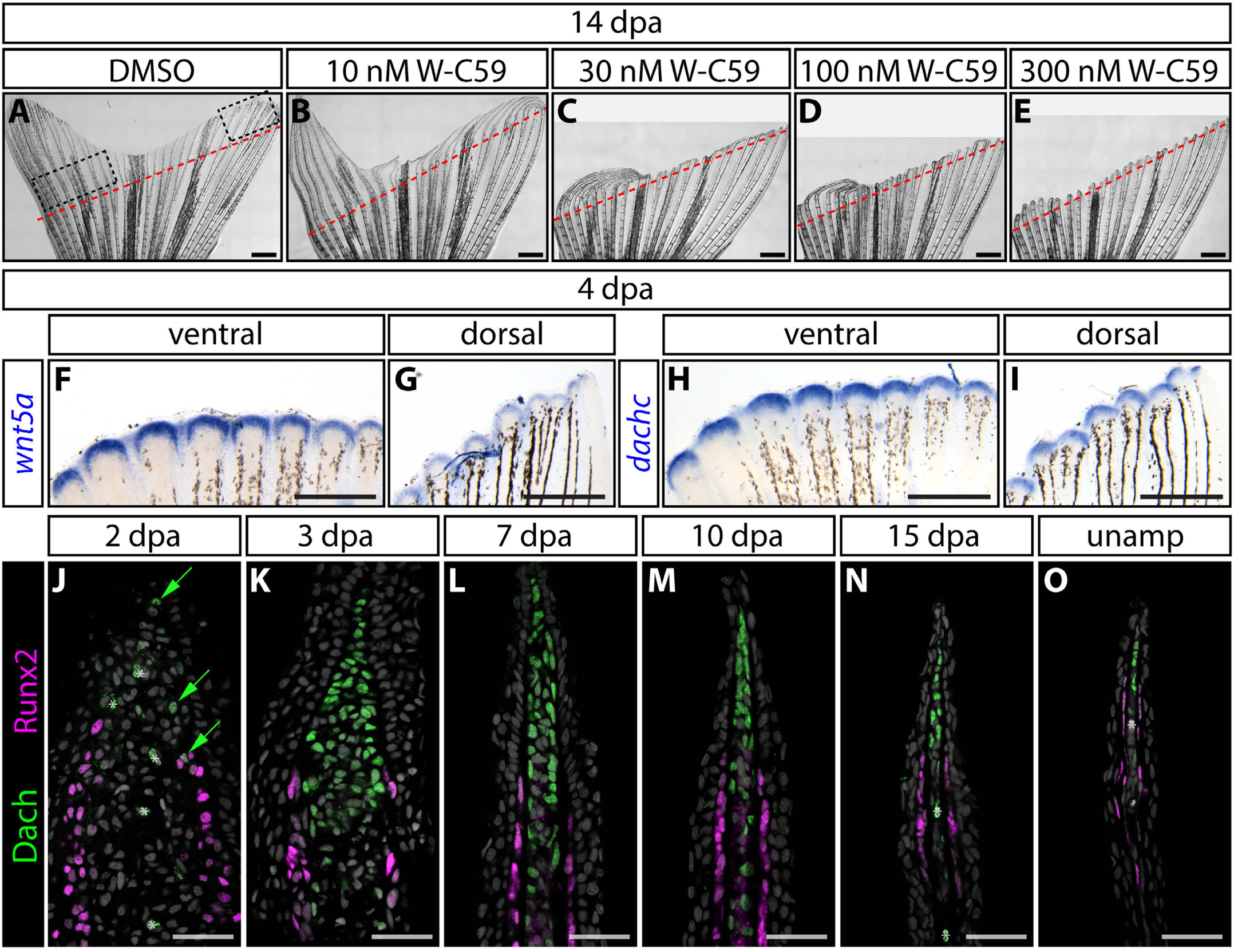
Dachshund transcription factors define a progressively depleting pool of distal niche cells in regenerating fins. (**A-E**) Sensitivity to Wnt inhibition inversely correlates with demand for regenerative growth. Diagonally fin-amputated zebrafish exposed to increasing concentrations of Wnt-C59 Porcupine inhibitor and imaged at 14 days post amputation (dpa). Red dashed lines show the amputation plane. Scale bars are 1 mm. (**F-I**) The size of distal *wnt5a* and *dachc* expression domains positively correlate with regenerative demand following diagonal amputations. Images compare whole mount RNA in situ hybridization signal intensities between ventral (proximally amputated) and dorsal (distally amputated) sites along the diagonal resection plane, approximated by dashed boxes in (A). Scale bars are 500 μm. (**J-O**) Dachshund (Dach) expression defines “niche” cells in the distal blastema, distinct from adjacent Runx2-expressing progenitor osteoblasts. Dach^+^ cells are first observed at 2 dpa, peak in number around 3 dpa, and then progressively decrease to a small residual pool that is also present in unamputated fins. Images show antibody-stained regenerating fin sections at the indicated times post-amputation. Dach and Runx2 are green and magenta, respectively. Hoechst-stained nuclei are grey. Asterisks denote red blood cells. Scale bars are 25 μm.

Wnt signaling “strength” could be encoded either by the amount of Wnt produced per distal blastema cell or the number of Wnt-producing cells. Regardless, we reasoned that understanding the properties of this unique population was key to deciphering size control mechanisms. Hereafter, we term these “niche cells” for brevity and because they generate the pro-progenitor and pro-growth environment that enables tissue expansion (Morrison and Spradling, 2008). We used RNA-Seq of 4 days post amputation (dpa) distal vs. proximal fin regenerates, as demarcated by the distal epidermal marker Tg(*shha:EGFP)* (Ertzer et al., 2007), to identify niche-characterizing factors (Fig. S3). Among transcription factors, the *dachshund* (*dach*) family members (Mardon et al., 1994; Shen and Mardon, 1997) *dacha* (5.9-fold) and *dachc* (6.0-fold) were notably distally enriched (Fig. S3). We also used the RNA-Seq dataset to confirm the distal enrichment of several Wnts, with particularly high levels of *wnt5a* and *wnt5b* (Fig. S3). *dachc* mimicked *wnt5a* in having notably higher expression levels in 4 dpa proximal tissue of diagonally amputated regenerating fins (Fig. 1H, I).

We used antibody staining to conclusively identify caudal fin cells that express Dacha/c (hereafter referred to as Dach) over the course of fin regeneration (Fig. 1J-O). Dach-expressing cells were rarely observed before 3 dpa (Fig. 1J). At 3 dpa, Dach became robustly expressed in distal cells adjacent to, but distinct from, Runx2^+^ pre-osteoblasts (Fig. 1K). Dach^+^ niche cells then steadily reduced over the course of regeneration to a small residual population also found in distal tissue of unamputated fins (Fig. 1L-O). We conclude Dach transcription factors mark Wnt-producing niche cells throughout the course of regeneration. The three-day delay prior to the appearance of Dach^+^ niche corresponds with the view that fin regeneration proceeds by an acute injury repair phase followed by a prolonged period of progressive outgrowth (Wehner and Weidinger, 2015). Finally, differential numbers of niche cells produced upon injury and then maintained through regeneration likely accounts for our observation that variable Wnt strength correlates with extent of fin outgrowth.

Wnt/β-catenin signaling is active in niche cells where it indirectly promotes further proximal mesenchyme proliferation (Wehner et al., 2014). However, a potential role for Wnt in maintaining the niche pool itself is unexplored. Using the new Dach marker, we found that pan-Wnt inhibition initiated at 4 dpa using Wnt-C59 depleted niche cells (Fig. S4). Wnt inhibition prevented outgrowth but did not disrupt osteoblast differentiation, joint formation, or skeletal maturation (Fig. S4), reinforcing our previous insight that Wnt/β-catenin promotes bone progenitor maintenance upstream of and in opposition to differentiation (Stewart et al., 2014). Wnt, whether canonical or non-canonical, appears to have an analogous autocrine role – maintaining the niche population in a state whereby it can drive continued regeneration.

The loss of niche cells upon Wnt inhibition allowed us to test if artificial depletion of the niche pool mimics normal termination of regeneration – as supported by our observation that Dach^+^ niche cells progressively decrease over regeneration. Wnt inhibition irreversibly blocked fin regeneration, with only rare outgrowth seen long after drug removal (Fig. S4). In contrast, re-amputating Wnt-inhibited fins re-initiated full regeneration. Small molecule inhibition of FGF receptor signaling prevented outgrowth to the same degree as Wnt inhibition (Fig. S4). However, as FGF inhibition did not deplete Dach^+^ niche cells (Fig. S4), regeneration resumed following drug washout, matching previous observations (Lee et al., 2005). Therefore, mitogenic FGF, which appears niche-expressed (Lee et al., 2005; Poss et al., 2000; Shibata et al., 2016) and is Wnt-dependent (Wehner et al., 2014), acts downstream of Wnt’s niche maintenance role. We propose niche cell numbers set dynamic levels of both signaling proteins and therefore the outgrowth rate. Regeneration ceases when the niche pool, and correspondingly Wnt and FGF production, depletes below an effective level. By this model, understanding instructive growth control mechanisms requires revealing niche cell origins and fates.

Regenerated fin tissues are derived from progenitor cells formed by partial de-differentiation of mature cells extant at the injury site (Wehner and Weidinger, 2015). We sought to determine which pre-existing cell type(s) generates Dach^+^ niche cells. An obvious candidate was intra-ray mesenchyme, including joint-associated fibroblasts, that are activated some distance proximal to the amputation site and express *tryptophan hydroxylase 1b* (*tph1b*) (Tornini et al., 2016). These cells, which contribute to the distal blastema, are a source of replacement ray mesenchyme (Tornini et al., 2016; Tornini et al., 2017). We used a mosaic lineage tracing system (Stewart and Stankunas, 2012) to permanently label intra-ray mesenchyme of individual rays in uninjured fins by transgenic mCherry expression (Fig. 2A-D). We then amputated these fins and double stained 4 dpa sections containing labeled rays and derived regenerating tissue with Dach and mCherry antibodies. Distal but not proximal mCherry-expressing blastema cells co-expressed Dach. Further, Dach^+^ niche cells largely co-expressed *Tg(tph1b:mCherry)* (Fig. 2E, F). Finally, we found the fin regenerate-expressed Msx transcription factor (Smith et al., 2008) marks both proximal blastema cells and distal Dach-expressing cells (Fig. 2G). We conclude Dach^+^ niche cells derive from intra-ray mesenchyme that initially populates the distal portion of the blastema.

**Figure 2.**
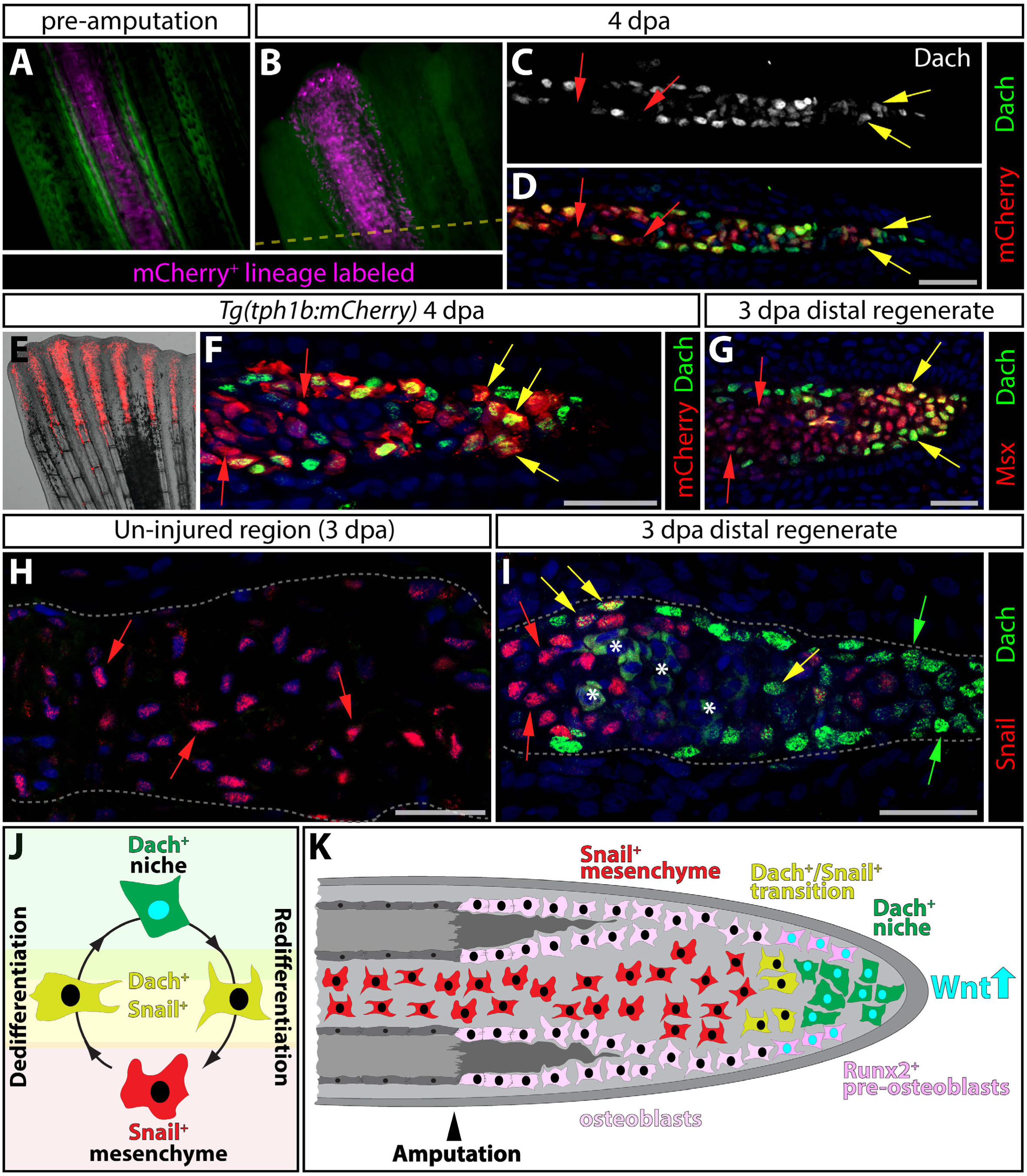
Niche cells are generated from state-transitioning intra-ray mesenchyme. **(A-D)** Genetic mosaic analysis demonstrates that the Wnt-producing niche derives from intra-ray mesenchyme. Example of a genetic mosaic fin with a single ray containing mCherry-labeled mesenchyme (in magenta) prior to amputation (A) and 4 dpa (B). The amputation plane is indicated with a dashed yellow line. (C, D) A section of the same ray at 4 dpa stained with anti-Dach (grey in single channel image in C, green in D) and anti-mCherry antibodies (red). Red arrows mark mCherry^+^ but Dach^-^ blastema cells and yellow arrows highlight Dach^+^/mCherry^+^ niche cells. **(E, F)** *Tg(tph1b:mCherry)* expression highlights the regenerating intra-ray mesenchyme/niche lineage. Immunostaining of a 4 dpa *Tg(tph1b:mCherry)* fin section shows Dach (green) and *tph1b* (red) expressing niche cells (yellow arrows). Red arrows mark further proximal Dach^-^/mCherry^+^ blastema cells. **(G-I)** Mesenchymal-to-niche cell transitions at 4 dpa defined by Dach, Msx, and Snail transcription factors. (G) Like *tph1b:mCherry*, Msx expression (red) labels the lineage whether in a differentiated mesenchyme (red arrows) or Dach-expressing (green) niche (yellow arrows) state. (H, I) Snail expression (red nuclei shown with red arrows) marks intra-ray mesenchyme proximal to the amputation site and blastema cells except distal Dach^+^ niche cells (green nuclei highlighted by green arrows). A small number of Snail^+^/Dach^+^ cells (yellow nuclei indicated by yellow arrows) identify cells transitioning between mesenchyme and niche states. Scale bars in all panels are 25 μm. Asterisks denote red blood cells. **(J)**. Model of cell state transitions between intra-ray mesenchyme and niche cells during regeneration. Snail^+^ mesenchyme cells normally populate the intra-ray space. After amputation, Snail^+^ cells dedifferentiate into Dach^+^ niche cells. As ray outgrowth proceeds, niche cells redifferentiate to contribute Snail^+^ intra-ray mesenchyme. **(K)** Control of regeneration by a Wnt-producing dynamic niche pool. Runx2^+^ osteoblasts (in pink) are maintained by Wnt (represented by cyan nuclei) secreted by the distal Dach^+^/Wnt^+^ niche (green cells with cyan nuclei), which transition (yellow cells) back to Snail^+^ mesenchyme (red cells) as regeneration progresses.

The mesenchymal cell state marker Snail (Lamouille et al., 2014) was expressed in un-injured intra-ray and proximal regenerate mesenchymal cells (Fig. 2H, I). In contrast, Snail was absent from Dach^+^ niche cells except for a small population of transitioning double Dach^+^/Snail^+^ cells (Fig. 2I). Concordantly, *snai2* transcripts were abundant in blastema mesenchyme but undetectable in distal-most tissue (Fig. S5). Dach-positive cells incorporated EdU even with a brief 4-hour pulse prior to tissue collection, showing the niche population includes actively proliferating cells (Fig. S5). These observations support a cyclical model whereby fin amputation induces a delayed transition of Snail^+^ intra-ray mesenchyme to a Dach^+^ niche state (Fig. 2J, K; Fig. S6). The Dach-expressing niche cells proliferate but, on net, progressively deplete through their conversion back to Snail^+^ mesenchyme that re-populates regenerated bony rays.

The origin of niche cells from intra-ray mesenchyme led us to consider that skeletal geometry might instruct how much niche is formed upon amputation. By this model, resecting a larger bony ray, with a relatively high volumetric capacity for mesenchyme, would produce a larger initial niche population and therefore more regeneration. In support, the size of fin ray bone segments noticeably tapers along the proximal-to-distal axis (Fig. 3A-E). Further, longer peripheral rays are wider at their base than shorter central rays. 3-D micro-CT analysis confirmed this assessment and underscored that the internal geometry of the rays approximates cylinders (Fig. S7; Supplemental Video 1). Therefore, we considered the activated stretch of a ray extending proximally from an injury site (Knopf et al., 2011; Tornini et al., 2016) as a cylinder of fixed length but variable width. As such, a ray’s mesenchyme-holding capacity, injury-induced niche size, and finally fin outgrowth should correlate with ray widths at injury positions. We measured the width of the central sixteen rays at a fixed proximal-distal position and then plotted the radius^2 values as adjacent bars. The bars’ distribution approximated that of ray lengths, largely matching the overall shape of a zebrafish fin (Fig. 3A).

**Figure 3.**
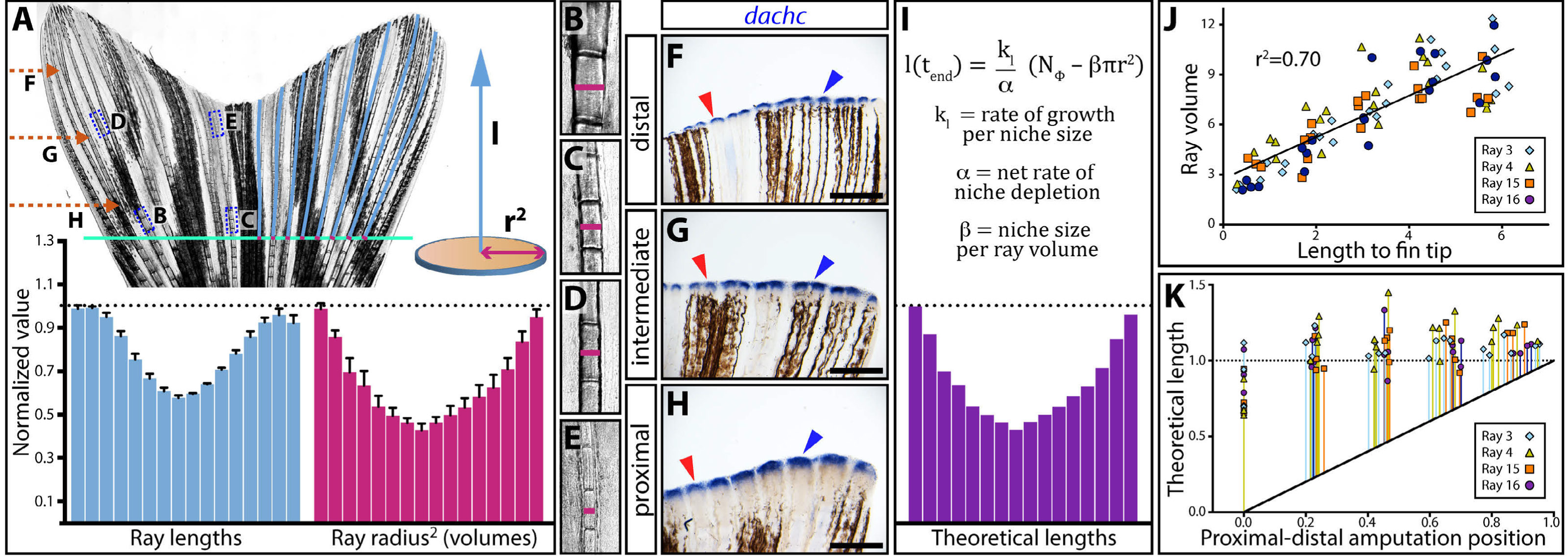
Skeletal geometry-dependent niche generation with progressive niche depletion models regenerated ray length. **(A)** A ray’s cross-sectional area (therefore, volume) exposed upon amputation (modeled by the square of the radius, *r^2^*, pink) anticipates the length (*l*, light blue) it will regenerate. Plotted normalized measurements for the sixteen central rays show a tight correlation between ray lengths and ray volumes, from/at a hypothetical proximal amputation position (green line). Error bars are one standard deviation (n = 4). (**B-E**) Scale-matched zoom images of the ray segments boxed in the stitched whole fin image in (A), showing differential girth depending on bone position. Pink lines mark ray diameters. **(F-H)** *dachc* whole mount in situ hybridizations on 72 hpa regenerating fins amputated at distal (F), intermediate (G), or proximal (H) positions. As anticipated by the model, *dachc* expression, representing the amount of niche generated, correlates with regenerative demand. Scale bars are 500 μm. **(I)** A mathematical model equation represents the length of a given ray at the end of regeneration (*l(t_end_)*) as a function of the radius of the ray at the amputation position. Three scaling parameters (*k_l_*, *α*, and *β*) must be tuned precisely to restore the original length, but no molecular positional information is required. The plot shows how the formula theoretically restores normalized ray lengths using actual measured radius values at the hypothetical amputation point (green line). **(J)** Scatter plot showing a linear relationship between ray volume and ray length from any measured position along the proximal-distal (P-D) fin axis. Each point represents 5 or 6 measured points from each of rays 3, 4, 15, 16 from four animals. Matching data point colors/shapes indicates a given ray. Units are arbitrary. **(K)** The mathematical model predicts regenerated length (normalized = 1) regardless of amputation position (normalized units). Each data point shows the theoretical length derived from a given ray’s radius measurements at a single P-D position. The colored vertical lines represent anticipated growth from the amputation position (therefore, starting from the black diagonal line).

This model predicts niche amount should vary by proximal-distal amputation position, correlating with the tapering bony rays. Whole mount in situ hybridization for *dachc* transcripts at 3 dpa, the time of niche formation, revealed that proximal amputations produced large *dachc*-expressing niche pools while distal cuts generated progressively smaller niches (Fig. 3F-H). Further, *dachc* expression across the dorsal-to-ventral fin axis correlated with regenerative demand upon perpendicular resections (Fig. S8) – as with diagonal amputations (Fig. 1H, I). Peripheral regions that re-form the longest rays expressed high levels of *dachc*. In contrast, central tissue that generates short rays to restore the original V-shaped fin produced small *dachc*-marked niches.

We used mathematical modeling to further explore if skeletal geometry coupled with progressive niche depletion could account for robust size restoration during fin regeneration. We first assumed outgrowth rate is proportional to the current number of distal niche cells. We considered the volume of intra-ray mesenchyme activated proximally to an amputation site (Tornini et al., 2016) determines the size of the starting niche pool. As validated earlier, ray radius^2 then serves as a proxy for the number of initial niche cells. A niche that collectively re-differentiates faster than it proliferates over the course of regeneration establishes a “countdown timer”. The number of initial niche cells sets the timer, with outgrowth ending when the niche population depletes below some effective level. As such, bony ray size at the amputation site ultimately instructs regenerated ray length.

We solved differential equations to derive a final equation describing regenerated ray length as a function of ray radius at the amputation position and three “scaling parameters”:

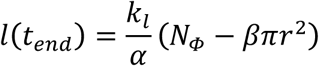

where *k_/_* is the rate of length growth per niche cell, *α* is the difference in niche cell growth rate versus rate of differentiation back to intra-ray mesenchymal cells, *N_Φ_* is the number of niche cells below which growth stops, and *β* is the number of niche cells released per injury-activated ray volume. Mathematically exploring the formula reinforced the conclusion that *α* must be negative to provide net niche depletion. Solving expected regenerated ray lengths using measured ray radii and optimized constants produced a distribution largely corresponding with actual measured lengths (Fig. 3I) – naturally matching normalized radius^2 distributions. A largely linear relationship between ray radius^2 and remaining ray length from the measurement position held across the proximal-distal fin axis (Fig. 3J, R-squared = 0.7). Likewise, existent ray lengths from a hypothetical injury point matched theoretical regenerated lengths derived from radius measurements and our growth equation (Fig. 3K).

Several zebrafish mutants develop and regenerate long fins, including dominant *longfin^t2^* (Van Eeden et al., 1996), well known as one of two mutations characterizing the *Tüpfel longfin* (*TL*) strain (Haffter et al., 1996) (Fig. 4A, B). *longfin^t2^* ray volumes estimated by normalized radius^2 measurements predicted overall fin shape but greatly under-anticipated ray lengths (Fig. 4C). Therefore, the skeleton was not proportionally re-scaled during development and instead suggesting *longfin^t2^* disrupts an outgrowth regulatory mechanism. We used our mathematical model to predict the growth-determining scaling parameter disrupted in *longfin^t2^* fish. The cell proliferation rate (*k_l_*) could be larger (e.g. increased mitogen production or mitogen sensitivity), *β* could be larger, establishing a larger niche pool upon injury, or *α* could be “less negative”, leading to niche perdurance. Changing *k_l_* or *β* vs. *α* had distinct theoretical effects when plotting expected regenerative growth over time and growth rate over time (Fig. 4D, E). Following amputation, *longfin^t2^* fish showed prolonged regenerative growth rather than increased initial growth or peak growth rate (Fig. 4F, G; Fig. S9), suggesting they have a persistent niche (less negative *α*) rather than enhanced niche generation (*β*) or increased cellular growth rate (*k_l_*).

**Figure 4.**
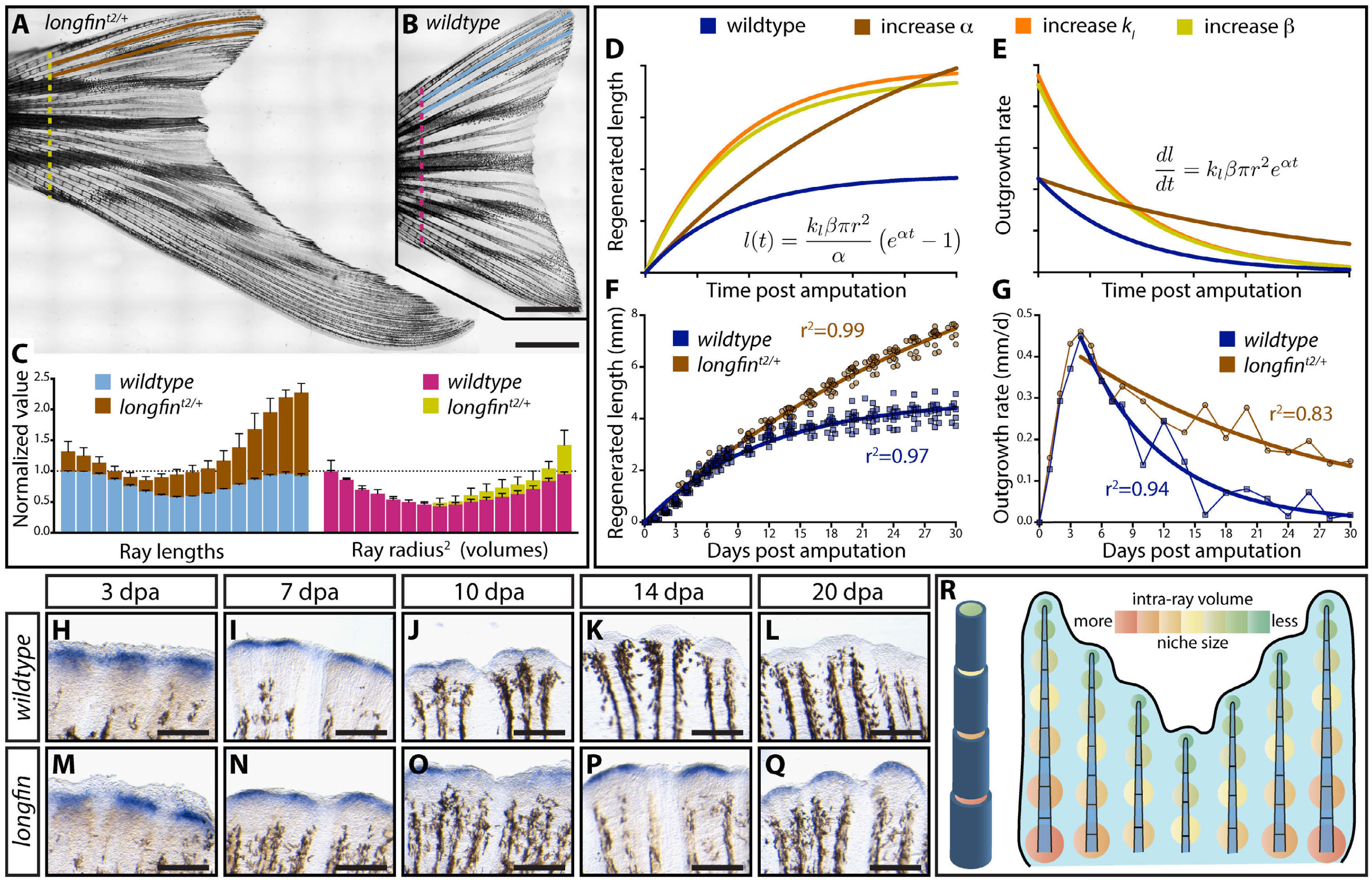
*longfin* fish regenerate exceptionally long fins due to a broken niche countdown timer, supporting a transpositional scaling model of self-restoring fin geometry. **(A, B)** Stitched differential interference contrast images show the dramatically overgrown caudal fins of *longfin^t2/+^*fish. Ray length and radius measurements taken at/from a proximal position denoted by colored dashed lines (brown/yellow=*longfin^t2/+^*, light blue/pink=wildtype). Scale bars are 2 mm. **(C)** Normalized caudal fin ray lengths (left) and volumes (from *radius^2^* measurements, right) for clutchmate wildtype and *longfin^t2/+^* fish. **(D)** Theoretical regeneration growth and **(E)** growth rate curves derived from the transpositional scaling equation have distinct shapes depending if niche response (*k_l_*) or niche generation (*β*) vs. niche perdurance (*α*) parameters are increased. The actual growth **(F)** and growth rate **(G)** of regenerating *longfin^t2/+^* caudal fins match expectations if niche perdurance (*α*) – the “countdown timer” – is disrupted. The initial and maximum growth rate is unaltered but the growth rate progressively decreases at a much slower rate. Curves show actual data fit to the modeling equations (r-squared goodness of fit values are shown). All data points (≥ 12 fish per time) are shown in (F); points in (G) are mean values. (**H-Q**) Whole mount in situ hybridization for the *dachc* niche marker through a time course post-caudal fin amputation for wildtype and *longfin^t2/+^* fish. *longfin^t2/+^* fish generate a normal-sized niche after fin resection but the cells fail to deplete in a timely manner. Scale bars are 200 μm. **(R)** A “Transpositional Scaling” model can explain how positional information stored in skeletal geometry combined with a state transitioning and depleting niche population directs a fin to regenerate back to its original size and shape. A wider amputated ray produces more niche cells from its intra-ray mesenchyme (A tapered, segmented ray is shown on the left, with intra-ray volume graded from orange (more) to green (less)). Niche cells deplete at a constant rate through their re-differentiation to mesenchyme (starting and then diminishing niche pools represented by same orange-to-green gradation as well as circle sizes on the right fin graphic). The diminishing niches maintain progressively smaller bone progenitor pools, regenerating tapered rays and inherently restoring skeletal geometry primed for future injury responses.

We examined *dachc* expression in *longfin^t2^* heterozygotes following fin resection to see if niche dynamics were disrupted as anticipated by our modeling (Fig. 4H-Q). *longfin^t2^* fish initially formed a normal-sized niche. However, unlike wildtype clutchmates, their *dachc*-expressing niche only slowly depleted, implicating a less negative *α* constant. Increasing *α* while inputting *longfin^t2^* ray radii measurements largely predicted actual *longfin^t2^* regenerated ray lengths (Fig. S10). These results support the robustness of the model and show that *longfin^t2^* disrupts the niche depletion system.

We conclude ray volumes (set by their widths) are “transposed” into regenerated ray lengths, and hence fin form, via the amount of intra-ray mesenchyme exposed upon ray resection. The mesenchyme transitions into a concordantly sized outgrowth-promoting niche pool that then steadily depletes, ultimately returning the regenerated fin to its original size and shape. In 1929, Samuel Nabrit similarly concluded: “… *in* [*fin*] *regeneration the rate of growth and consequently the form is controlled locally by the cross-sectional area of the fin rays exposed*” (Nabrit, 1929). Therefore, Nabrit’s predicted growth promoting “fin ray end substance” likely is the injury-released intra-ray mesenchyme itself. Positional information stored in tapering ray skeletal geometry is self-restoring since a depleting niche progressively directs slower outgrowth and narrower bones due to its diminishing progenitor-maintenance capacity. We use the term “transpositional scaling” to describe this model for organ size determination – a new answer to an age-old question of appendage regeneration.

Vertebrate appendage regeneration, including that in zebrafish fins, is used as compelling evidence for pattern formation by Cartesian coordinate-defined cellular positional identities (Wolpert, 2016). However, our transpositional scaling model does not require fin cells retain any such molecular information. Similarly, the hypothesized “pre-pattern” established early in a regenerating blastema that directs regional outgrowth (Nachtrab et al., 2013; Rabinowitz et al., 2017; Rolland-Lagan et al., 2012; Tornini et al., 2016) can be explained by skeletal geometry-defined niche cell numbers alone. The state transition mechanisms that generate the niche and drive its progressive depletion are then identical throughout the fin but produce differential, predictable output depending on the starting condition defined by ray volumes.

Our conclusion linking *longfin^t2^* to disrupted niche depletion shows how a mutant can have profound effects on fin morphology by altering the α scaling parameter defined by our mathematical model. As such, we provide a framework to understand evolutionary origins of appendage size and shape variation. In the accompanying manuscript, we identify *cis* ectopic expression of the voltage-gated potassium channel *kcnh2a* in mesenchyme/niche lineage as the long sought cause of *longfin^t2^* (Stewart et al., 2019). These studies link ion signaling or bioelectricity, widely connected to organ size control and regeneration (McLaughlin and Levin, 2018), to gene regulatory pathways regulating cell state transitions that slow then terminate fin outgrowth.

Could “transpositional scaling” models apply in other circumstances where an organ is restored to form and scale? The hemi-ray bones in zebrafish fins appose to approximate cylinders, conveniently enabling consideration of the geometric storage of a growth-promoting cell lineage. Urodele amphibian limb regeneration is another striking model of size and pattern restoration. Matching our idea, an older model proposes that initial blastema mass rather than origin determines which proximal-distal limb skeletal elements regenerate (de Both, 1970). However, urodele limb cartilage/bones are solid and therefore position-defining geometric reservoirs of a hypothetical niche lineage would have to lie around rather than inside skeletal structures. Intriguingly, urodele limb positional information is stored in connective tissue mesenchyme surrounding bones, albeit reportedly by molecular memories (Kragl et al., 2009; McCusker et al., 2016; Nacu et al., 2013). Regardless, progenitor cell supporting-niche populations are a common component of organ growth, homeostasis, and regeneration (Chacón-Martínez et al., 2018; Yamashita and Tumbar, 2014). Our simple model, inherently dependent on regulatory networks with the positional information stored in tissue geometry rather than molecular information, establishes a compelling paradigm to explore in many contexts.

## MATERIALS AND METHODS

### Zebrafish

Wildtype AB (University of Oregon Zebrafish Facility), *longfin^t2^* (Haffter et al., 1996; Maderspacher and Nüsslein-Volhard, 2003; Van Eeden et al., 1996), *Tg(-2.4shha:gfp:ABC)* (Ertzer et al., 2007), *Tg(tph1b:mCherry)* (Tornini et al., 2016), *Tg(dusp6:CreERT)* (Stewart and Stankunas, 2012), and *Tg(eab:FlEx)* (Boniface et al., 2009) zebrafish lines were used. Zebrafish were housed in the University of Oregon Aquatic Animal Care Services facility at 28-29°C. The University of Oregon Institutional Animal Care and Use Committee oversaw animal use.

### Transgenic mosaic fin lineage tracing

Zebrafish embryos carrying the *Tg(dusp6:CreERT)* and *Tg(eab:FlEx)* lines were treated with low doses of tamoxifen, reared to adulthood, screened for mosaic labeling, caudal fin-amputated, and analyzed by whole mount imaging and antibody staining of sectioned tissue as described previously (Stewart and Stankunas, 2012).

### Small molecule studies of fin regeneration

The Porcupine inhibitor Wnt-C59 (Proffitt et al., 2013), which blocks Wnt secretion, and the FGF receptor tyrosine kinase inhibitor PD173074 (Mohammadi et al., 1998) were dissolved and diluted in DMSO. Caudal fins were amputated at a fixed proximal-distal position or common oblique plane (illustrated in Fig. S1). Fin-resected fish were then immediately placed in drug-containing aquarium water. Animals were transferred to fresh drug-containing water every 48 hours. The final concentration of DMSO in fish water was 0.01% for all conditions.

For niche depletion experiments with delayed small molecule addition (Fig. S4), fin-resected AB wildtype fish first were allowed to regenerate for 4 days. The animals then were treated with DMSO (0.01%, n = 10), Wnt-C59 (500 nM, n = 10), or PD173074 (5 μM, n = 10) for another 4 days. Animals were fed and changed to fresh drug-containing water after 48 hours. At 8 days post-amputation (dpa), animals were returned to plain fish water. In some cases, fins were subjected to a second amputation proximal to the original amputation site at 12 dpa. Animals were followed for up to 60 dpa, when each fish was imaged using a Leica M165 FC stereomicroscope. Representative fish for each condition were visualized using high resolution stitched imaging (Nikon Eclipse Ti widefield microscope and NIS-Elements software) before drug treatment, immediately after drug treatment, and at the experiment’s conclusion. Regeneration was quantified by scoring the number of fully regenerated individual rays. This complete experiment was repeated three times with the same outcomes.

### RNA-Seq

Caudal fins of adult *Tg(-2.4shha:gfp:ABC)* zebrafish were resected. At 4 dpa, dissected regenerated tissue at and beyond the distal GFP-marked epidermal domains was collected as “distal regenerate” samples. The remaining proximal regenerated tissue also was harvested. Tissue pooled from four animals constituted matched replicate samples. Tissue was immediately placed in TRIzol reagent (Thermo Fisher) and homogenized. RNA was isolated following the manufacturer’s instructions with minor alterations. The RNA was precipitated overnight at -80°C and then pelleted at 15,000 rpm for 30 minutes at 4°C. Pellets were washed twice with 70% ethanol, dried for 10 minutes at room temperature, and resuspended in RNase-free water (Thermo Fisher).

RNA-Seq libraries were prepared from 1 μg of isolated RNA using a Kapa Biosystems Stranded mRNA-Seq kit. Bar-coded libraries were pooled and sequenced using a NextSeq 500 (Illumina). Illumina reads were aligned to the zebrafish genome (GRCz11) using TopHat2 (Kim et al., 2013). Gene-assigned aligned reads were counted using HTseq (Anders et al., 2015). DEseq (Anders and Huber, 2010) was used to determine differential expression of 1147 genes annotated to have “transcription regulator activity” (GO:0140110) by AmiGO 2 (Ashburner et al., 2000; Carbon et al., 2009; Gene Ontology Consortium et al., 2013). Genes below the 40% quantile for total counts were filtered out to improve the statistical analysis of relatively highly expressed genes.

### Immunostaining and in situ hybridization

Immunostaining was performed on paraffin sections as described (Stewart et al., 2014). Antibodies used were: Runx2 (Santa Cruz Biotechnology, 27-K, 1:500-1000 dilution); Dach (Proteintech Group, 1:2000); Msx (Developmental Studies Hybridoma Bank, tissue culture supernatant, 1:15-20); Snail (Developmental Studies Hybridoma Bank, ascites, 1:100); mCherry (Takara Bio or Novus Biologicals, 1:100). For Snail/Dach double immunostaining, Dach antibodies were used at 1:1000 and sodium dodecyl sulfate (SDS) was added to 0.005% during the primary antibody incubation step. Immunostained sections were imaged using either Olympus FV1000 or Zeiss LSM 880 laser scanning confocal microscopes.

Whole mount in situ hybridization using DIG-labeled probes followed established procedures (Stewart et al., 2014). The *wnt5a* probe was described previously (Stewart et al., 2014). Templates for *dacha* and *dachc* probes were amplified by PCR from regenerating fin cDNA using the indicated primers, where the reverse primer contains a T7 promoter sequence (in caps) for in vitro transcription using DIG labeling mixtures (Roche) and T7 RNA polymerase:

*dacha* forward: 5’-atggccgtatctgcaactcctccggtgc-3’

*dacha* reverse: 5’-TAATACGACTCACTATAGGGtcagtacatgatggggggtttggagtagg-3’

*dachc* forward: 5’-atggcccacgcgcctccga-3’

*dachc* reverse: 5’-TAATACGACTCACTATAGGGtcagtacatcattgtggactttagaaagagcctt-3’

The *snai2* probe template was generated by PCR amplifying its coding sequence from regenerating fin cDNA using the following primers followed by ligation to the pCRII vector (Thermo Fisher).

*snai2* forward: 5’-atgcctcgttcattcctagtaaagaagc-3’

*snai2* reverse: 5’-tcagtgtgcgatgcaacagccag-3’

The resulting plasmid was linearized and a DIG-labeled probe synthesized by in vitro transcription using SP6 RNA polymerase.

### 5-ethynyl-2’-deoxyuridine (EdU) incorporation and staining

10 μg of EdU in saline was intraperitoneally injected into wildtype fish 92 hours post caudal fin amputation. Fin tissue was collected 4 hours later, fixed, and processed for paraffin sectioning. EdU detection used a Click-iT kit (Thermo Fisher) combined with immunostaining as described above using Dach antibodies.

### X-ray micro computed-tomography imaging

An adult wildtype zebrafish was sacrificed and fixed in 4% PFA for 24 hours. The specimen was stabilized within a 15 ml conical tube and its caudal fin scanned using a VivaCT 80 (SCANCO Medical) with 55 kVp X-ray source energy, 145 µA current, 6.5 µm pixel size and 1000 ms per projection integration time. Slice images were reconstructed to a 1590x1590 pixel matrix using an automated cone beam convolution backprojection algorithm. The resulting 2-D images were output for 3-D reconstruction using Imaris 9.3 software (Bitplane).

### Morphometrics and mathematical modeling

Rays were considered as simple cylinders to facilitate modeling regenerative fin outgrowth as a function of skeletal geometry. A given bony ray segment’s volumetric capacity for injury-activated, niche-forming intra-ray mesenchyme (Tornini et al., 2016), then could be estimated by first measuring the ray’s width (= diameter) at a hypothetical amputation site. For modeling purposes, the square of the ray’s radius was used as a proxy as normalized areas and volumes (for cylinders of a fixed length) are mathematically identical.

Caudal fins of five-month-old *longfin^t2/+^*and wildtype clutchmate fish were imaged using differential interference contrast microscopy and multi-field stitching (Nikon Eclipse Ti and NIS-Elements) for morphometrics. Fish were euthanized by tricaine bath overdose and mounted in 0.75% low melt agarose / PBS on microscope slides. The fins were gently fanned out using a horse hair brush to fully extend the tissue and separate rays. The initial proximal-distal positions for ray measurements were determined by first identifying the end of the innermost truncated rays (peripheral “mini-rays” that, unlike the standard 18 caudal rays, do not extend the full distance of the fin, shown in Fig. S1). A line was drawn across the fin that intersected points two ray bone segments preceding the end of the mini-rays on the fin’s dorsal and ventral sides. The width of each ray was measured where it intersected this transverse line.

Similarly, polylines (using anchored points to allow for shifts in ray orientation) starting from the same proximal-distal position were drawn along each ray and then directly in between branches of a given “mother ray” until reaching the fin tip. The lengths of these lines defined how much each ray would have to regenerate to restore the fin’s original size. Lengths were normalized to the longest wildtype ray. Repeated measures taken every three bone segments along the proximal-distal axis for rays 3, 4, 15, 16 used the same procedure, with lengths representing the distance from the diameter measurement position to the end of the ray.

We derived a mathematical model for “Transposition Scaling” that predicts the extent of regenerative outgrowth as a function of skeletal geometry – the bony ray’s capacity for niche-originating mesenchyme. First, we considered the number of niche cells released at a cut site as:

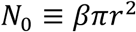

where *r* is the radius of the ray at the cut site and *β* is the average number of niche cells released per unit area. Niche cells can then proliferate (given by *k*_*g*_) or convert to a differentiated, mesenchymal state (given by *k*_*q*_). The change in the number of niche cells per unit time is:

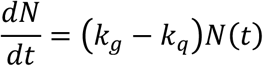

The difference in these rates can be summarized by α ≡ (*k*_*g*_ − *k*_*q*_). Using the boundary condition that *l*(0) = 0, the solution to this first-order differential equation is: 

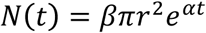

We model bony ray growth rate as directly proportional to the number of niche cells, controlled by *k*_*l*_:

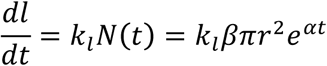

Using the boundary condition that *N*(0) = *N*_0_, the solution to this differential equation is:

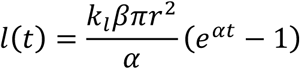

To link the number of cells released to the final ray length, we assume that skeletal growth stops when the number of cells reaches *N*_*Φ*_, the minimum number of cells required to support growth. This happens at *t*_*end*_:

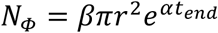

We then solve for *t*_*end*_:

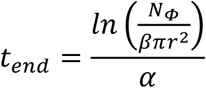

and substitute into our model for *l*(*t*):

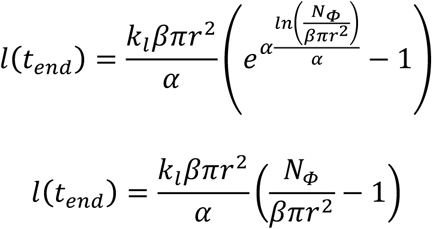

This gives the final relationship:

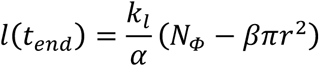

where *k*_*l*_ is the length growth rate per niche cell, α is the difference between the niche cell growth rate and their conversion back to intra-ray mesenchymal cells, *N*_*Φ*_ is the minimum number of cells required to support growth, and *β* reflects how many niche cells are released per bony ray cross-sectional area.

### Fin regeneration outgrowth measurements

Caudal fins from clutchmate *longfin^+/+^ and longfin^t2/+^* animals were amputated at the same position along the proximal-distal axis (Fig. S1) and allowed to regenerate for 30 days.

Outgrowth measurements were taken every 24 hours from 1-8 dpa and every 48 hours from 10-30 dpa. The length of the third ray (Fig. S1) from the amputation site to the fin’s tip was measured from images acquired with a Leica M205 FA stereomicroscope. Stitched fin images of zebrafish euthanized with tricaine and mounted in 0.75% low melting agarose were captured using a Nikon Eclipse Ti widefield microscope and NIS-Elements software. Data points were fit to user-defined transpositional scaling equations (see Mathematical modeling) by nonlinear regression using GraphPad Prism. The α, β, and *k_l_* parameters were allowed to vary to generate best-fit curves. *β* and *k_l_*were nearly identical between the wildtype and *longfin^t2/+^* data sets. In contrast, α was 3.0x higher for *longfin^t2/+^*, similar to the α difference (2.1x) derived from the geometrical (rather than growth rate) analysis.

## Supporting information

Supplementary RNA-Seq Analysis

## ACKNOWLEDGEMENTS

We thank C. Kimmel and V. Devasthali for feedback on the manuscript; the Stankunas lab for discussions; E. Shaw for technical assistance; the University of Oregon X-ray Imaging Core and A. Lin for assisting with micro-CT; K. Poss and ZIRC for providing zebrafish lines; and the DSHB for antibodies.

## FOOTNOTES

### Author contributions

S. S., G. A. Y., H. K. L. and K. S. designed experiments. S.S., G. A. Y., H. K. L., A. L. H., J. A. B., and J. C. performed experiments. K. S. and M. J. H. led the mathematical modeling. K. S. and S. S. prepared and wrote the manuscript with input from G. A. Y., H. K. L., M. J. H., and A. L. H.

### Data and material availability

RNA-Seq data are deposited at the NCBI Gene Expression Omnibus (GSE125859). Requests for materials should be addressed to S. S. and K. S.

---

**Supplemental Figure 1.**
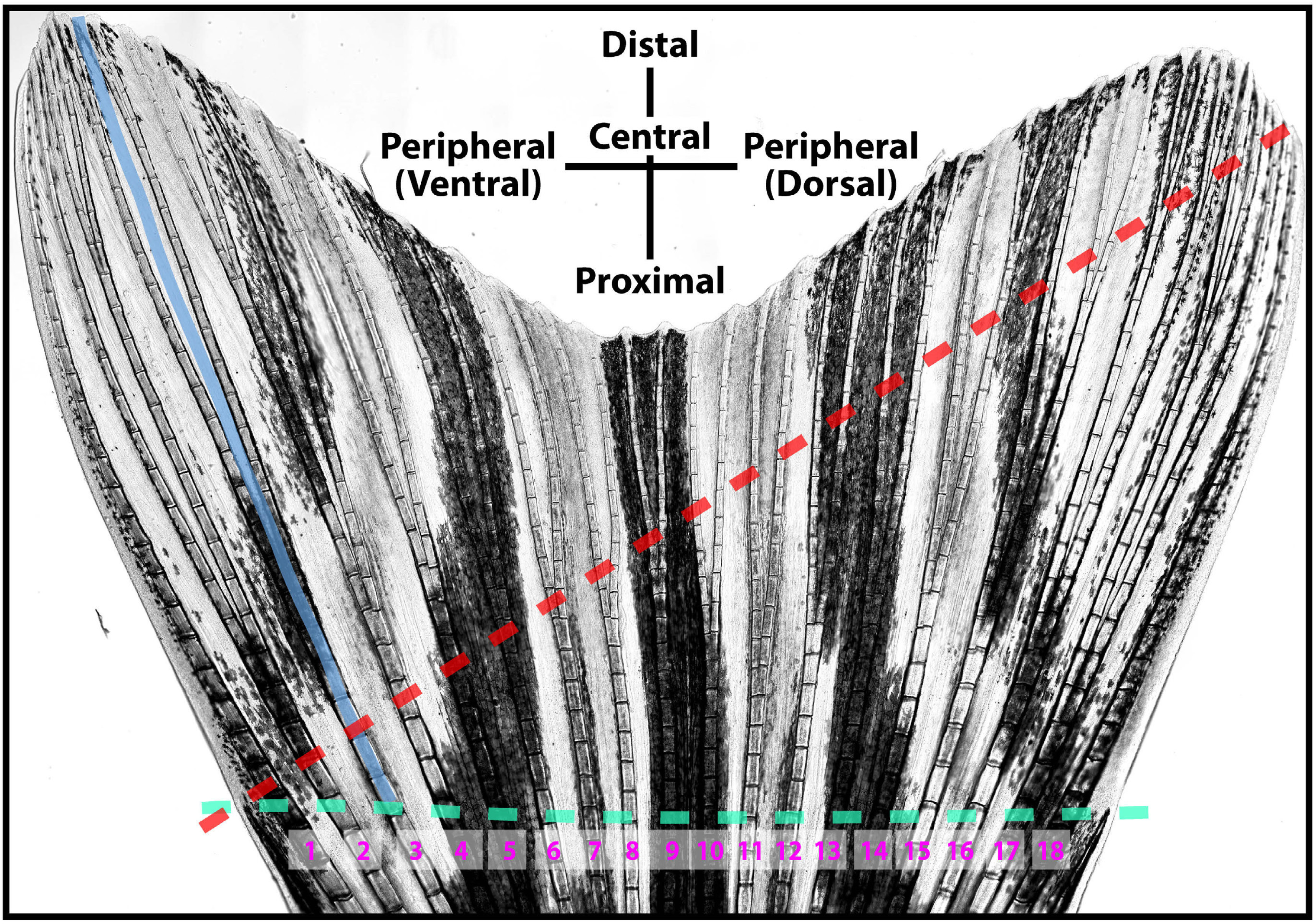
A zebrafish caudal fin showing amputation positions and orientation/numbering conventions. A high resolution, stitched differential interference contrast (DIC) image of an uninjured adult caudal fin. The dashed green line shows the standard proximal-distal amputation plane spanning between the tips of the un-numbered, truncated peripheral “mini-rays”. Diagonal fin amputations follow the dashed red line, starting from the tip of the ventral-most “mini-ray”, passing through ray 16’s second branch-point, and continuing straight through the dorsal edge. Length measurements (blue line) are from a real or theoretical amputation position to the fin’s tip. To accommodate branching, the line is drawn directly in between branches of a mother ray. The terms “central” and “peripheral” distinguish ray position relative to a fin’s center or either edge.

**Supplemental Figure 2.**
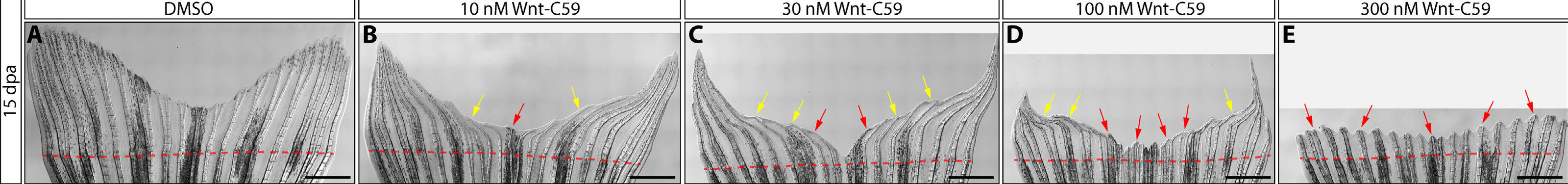
Wnt production correlates with regenerative “demand”. **(A-E)** Stitched differential interference contrast (DIC) images of regenerated caudal fins 15 days post amputation (dpa) with exposure to DMSO (A) or indicated concentrations of the Wnt-secretion inhibitor Wnt-C59 (B-E). Small molecule exposures began at 4 dpa until imaging at 15 dpa. Yellow and red arrows point to regions with reduced or failed regeneration after drug addition, respectively. The amputation plane is indicated with a dashed red line. Scale bars are 1 mm.

**Supplemental Figure 3.**
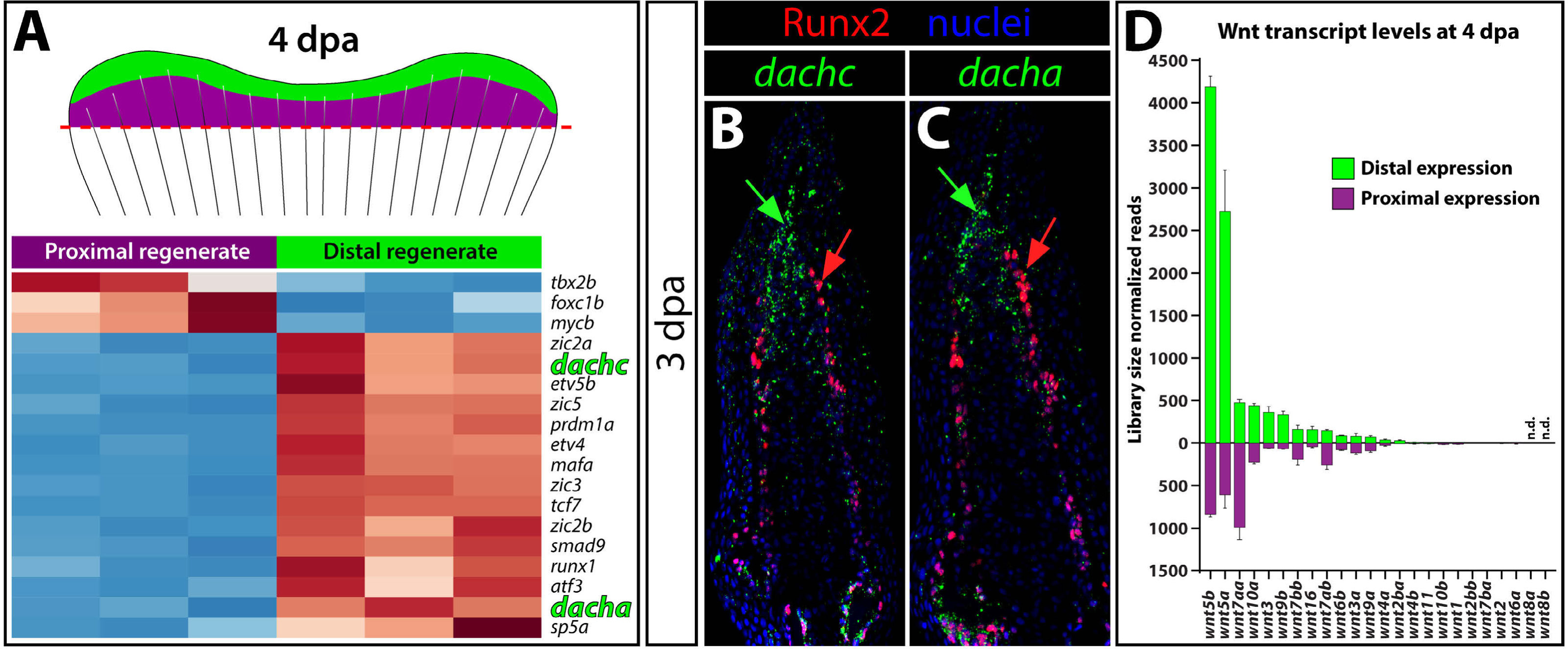
d*a*cha and *dachc* expression defines the distal Wnt-producing niche. **(A)** RNA-Seq analysis comparing distal Wnt-expressing vs. proximal differentiating tissue from 4 day post-amputation (dpa) regenerating fin tissue. The heat map illustrates relative enrichment (red = higher, blue = lower) of differentially expressed transcriptional regulators (p < 0.05 and at least 3-fold difference) for each replicate. **(B-C)** In situ hybridization (in green) for *dachc* (B) and *dacha* (C) mRNAs, combined with immunostaining for Runx2 (red) on 3 dpa fin sections. Hoechst-stained nuclei are blue. Red arrows indicate Runx2-expressing osteoblasts that lack *dacha/c* expression. Green arrows point to the *dacha/c*-expressing distal niche. **(D)** Bar graph showing the library size-normalized transcript counts of all 24 zebrafish *wnt* genes in distal and proximal 4 dpa fin regenerate tissue. The genes are ordered by descending expression levels. *wnt5b* and *wnt5a* have the highest distal transcript levels and are distally enriched. Means of the three replicate pooled samples are shown. Error bars show one standard deviation.

**Supplemental Figure 4.**
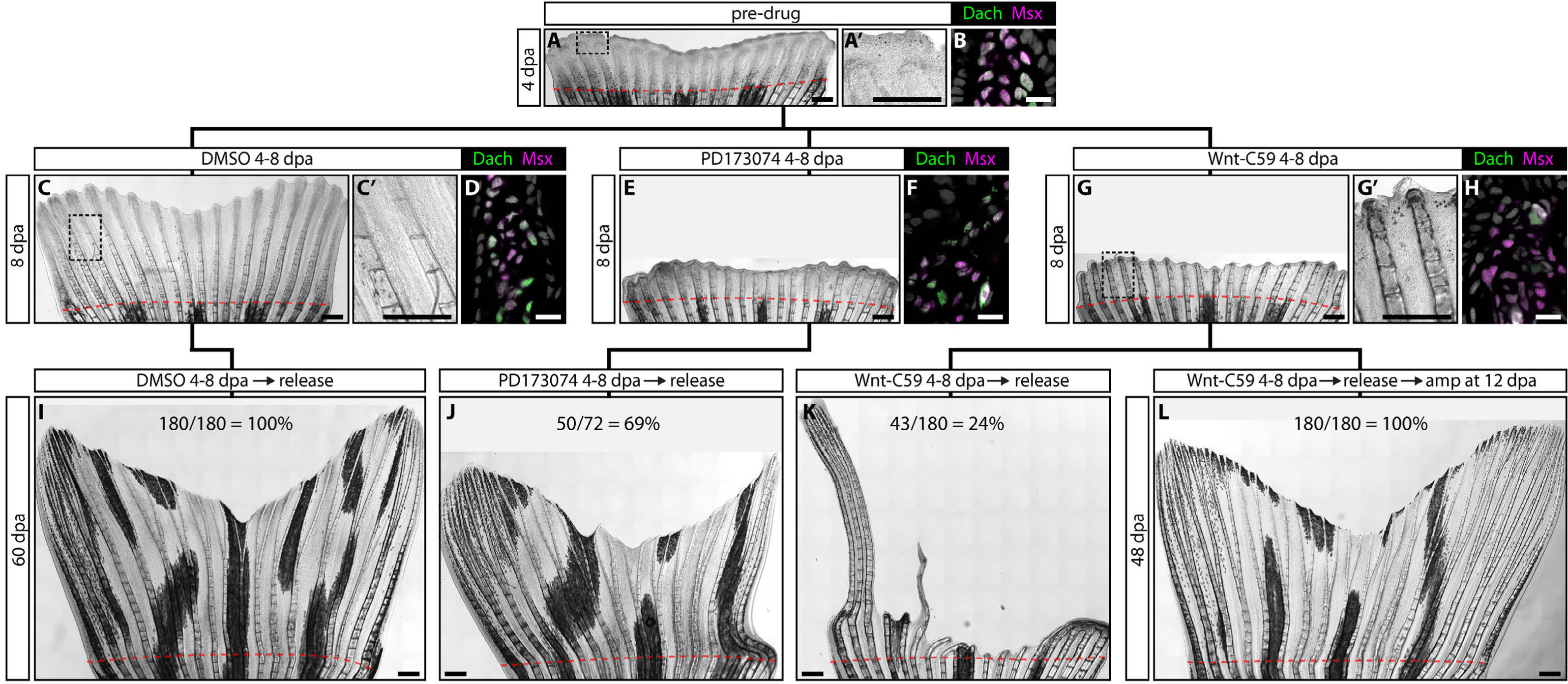
Inhibition of Wnt signaling depletes the Dach^+^ niche and irreversibly blocks regeneration. The black lines connecting figure panels establish a tree diagram showing the experimental workflow. Top row: caudal fins from clutchmate animals are resected and allowed to regenerate through 4 dpa. Middle row: At 4 dpa, animals are treated with DMSO, the FGFR inhibitor PD173074, or the Wnt secretion inhibitor Wnt-C59 until 8 dpa. Bottom row: Animals are then transferred to fish water and allowed to completely regenerate. For (L), fins are re-amputated at 12 dpa proximal to the original amputation site. Whole mount fin images use stitched differential interference contrast (DIC) microscopy. **(A)** A representative fin at 4 dpa. The region bound by the dashed box is shown at high magnification (A’) to emphasize how distal tissue is undifferentiated. **(B)** A distal regenerate section from a 4 dpa fin immunostained with Dach (green) and Msx (magenta) antibodies. Niche cells express both transcription factors. **(C, C’, E, G, G’)** 8 dpa regenerated fins from DMSO (C, C’), PD173074 (E), and Wnt-C59 (G, G’) exposed animals. FGF or Wnt inhibition prevents outgrowth past the 4 dpa time of drug addition. However, Wnt is not required for bone differentiation or joint formation (compare G’ with A’). (**D, F, H**) 8 dpa sections immunostained with Dach (green) and Msx (magenta) antibodies. Dach-expressing niche cells persist upon FGFR inhibition but are lost when Wnt production is blocked. **(I-K)** Fin images at 60 dpa after treatment from 4-8 dpa with DMSO (I), PD173074 (J), or Wnt-C59 (K). Unlike FGFR inhibition, Wnt inhibition irreversibly blocks outgrowth. **(L)** A Wnt-C59 treated animal (from 4-8 dpa) subjected to a secondary amputation at 12 dpa and allowed to regenerate until 48 dpa. A secondary amputation effectively “resets” the outgrowth timer by establishing a new niche, allowing regeneration to resume. For (I-L), the number of rays that regenerated normally is shown for each treatment (n = 10 fish for each condition). A dashed red line indicates the amputation plane. Scale bars are 500 μm for whole mount images and 25 μm for immunostained sections.

**Supplemental Figure 5.**
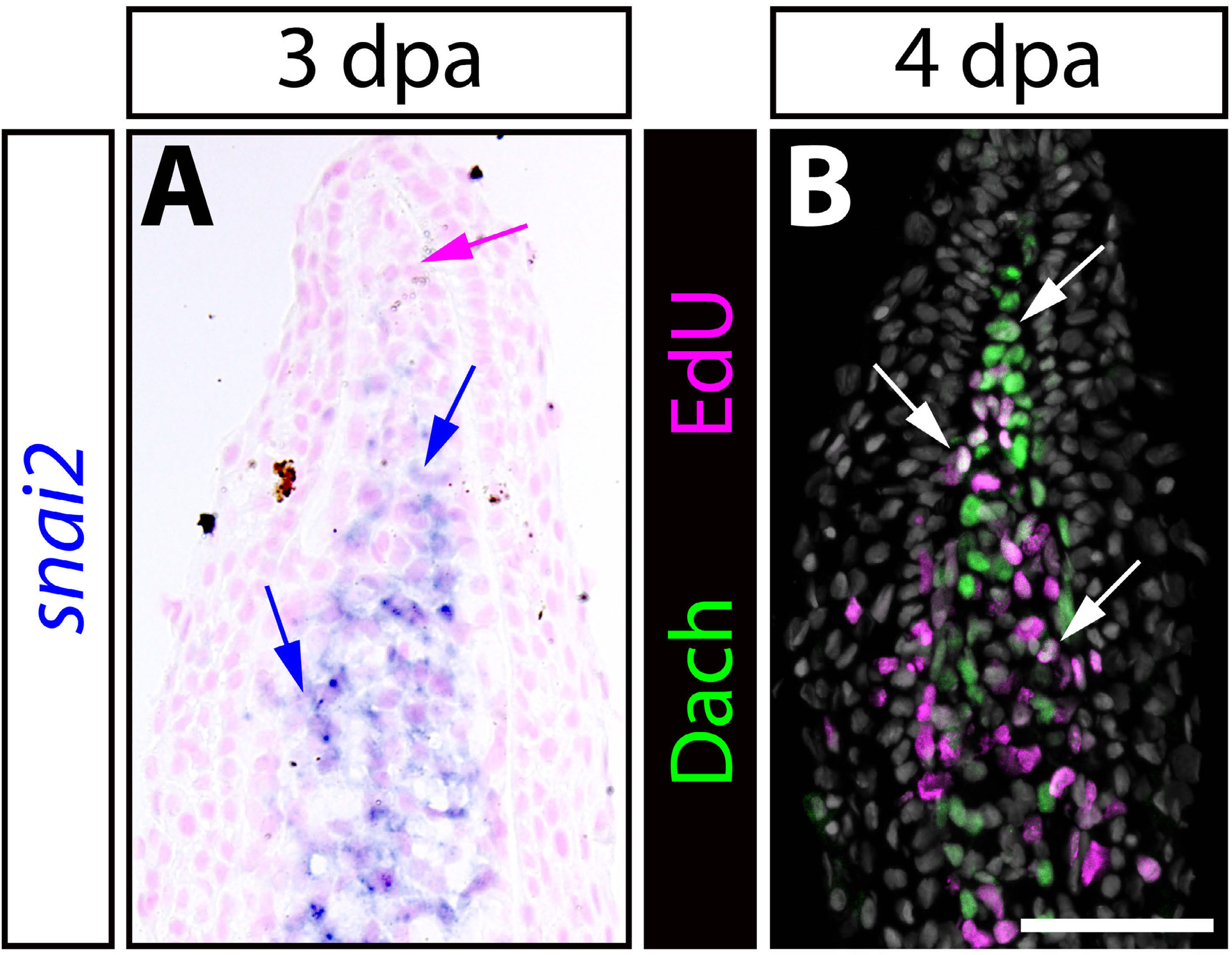
Intra-ray mesenchymal cells down-regulate *snai2* when transitioning to a Dach^+^ niche state that remains proliferative. **(A)** Visualization of *snai2* mRNA (in blue) by in situ hybridization on a 3 dpa fin section. The tissue is counterstained with Nuclear Fast Red. Blue arrows indicate *snai2*-expressing blastema mesenchyme. The magenta arrow highlights the *snai2*-negative niche. **(B)** Paraffin section of a 4 dpa regenerating caudal fin stained for EdU incorporation (four hour EdU exposure; magenta) and Dachshund (Dach; green) expression. Nuclei are in grey. Scale bar is 50 μm. White arrows denote Edu^+^/Dach^+^ cells found throughout the niche population.

**Supplemental Figure 6.**
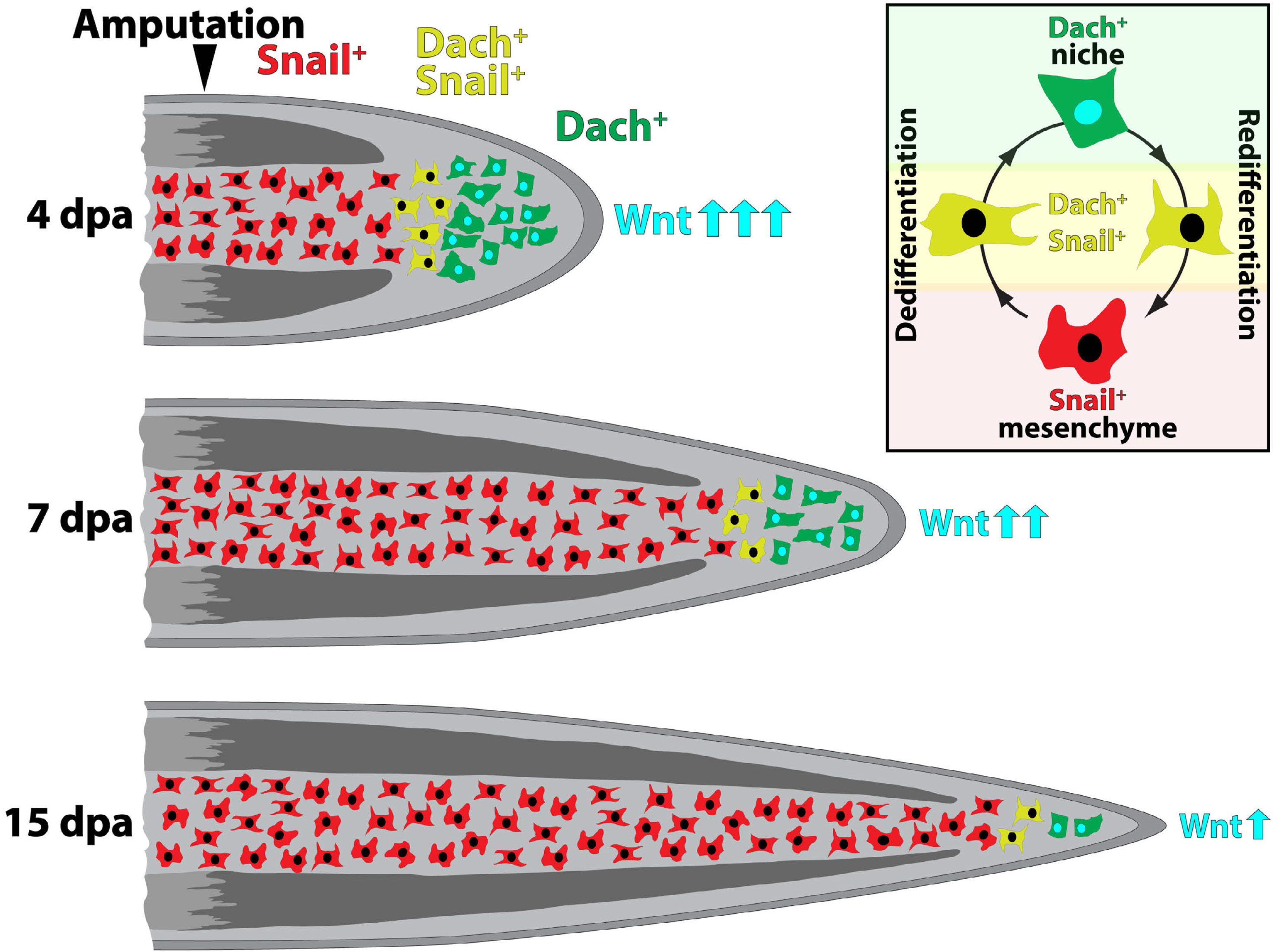
Graphic depicting the progressive depletion of the Wnt-producing, Dach-defined niche population during fin regeneration. Dach^+^ niche cells originate from Snail-expressing intra-ray mesenchyme released into the forming blastema upon injury. The size of the niche population, and therefore Wnt levels, decreases as the fin regenerates and niche cells transition back into a Snail^+^ mesenchymal state.

**Supplemental Figure 7.**
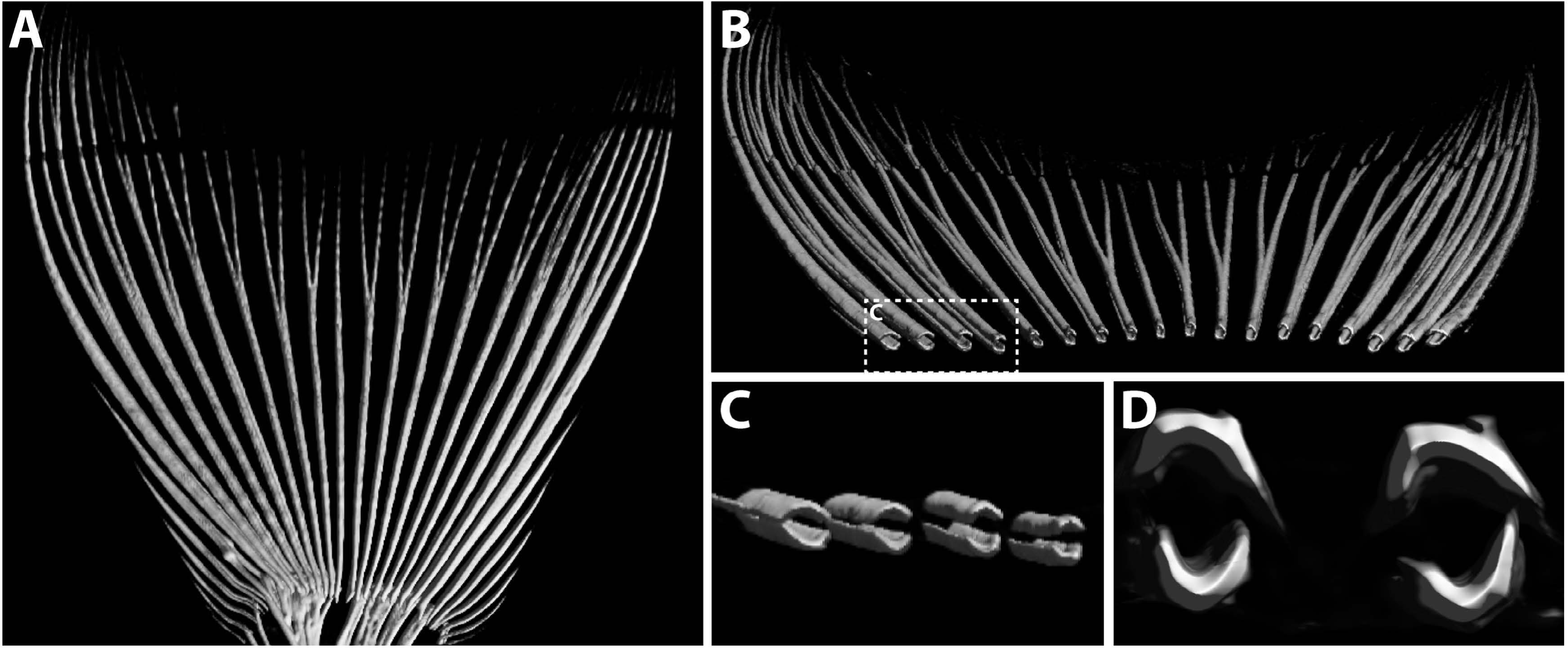
Micro-CT analysis shows bony rays across the caudal fin are variably sized, tapered, and cylindrical. **(A-C)** 3-D micro-computed tomography images of an adult zebrafish caudal fin highlighting its (A) bony ray skeleton, (B) differentially sized and tapering rays, (C) the cylindrical shape of the intra-ray space between hemi-rays and (D) its approximately circular cross-sectional area.

**Supplemental Figure 8.**
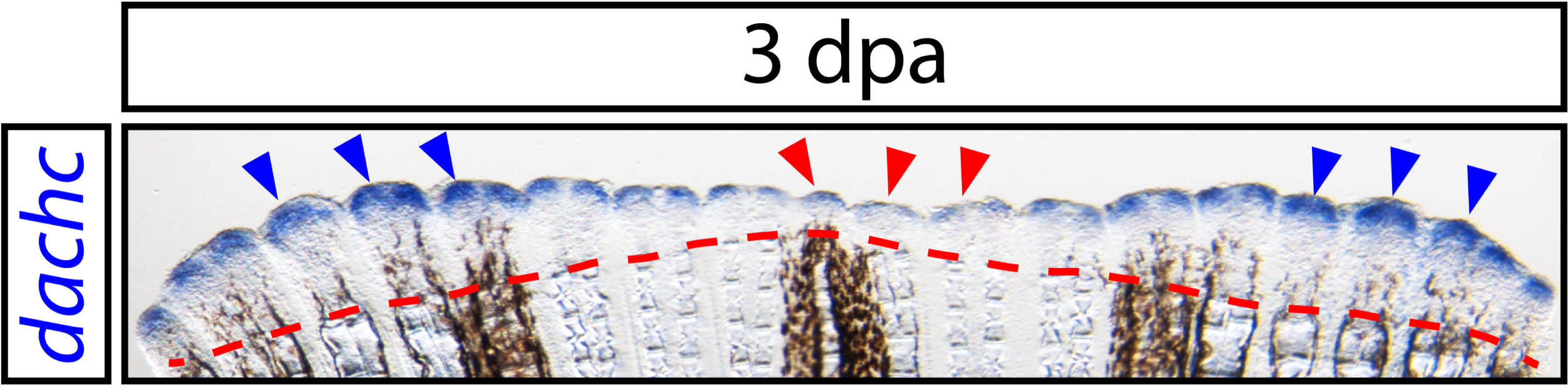
Differential niche production across the dorsal-ventral axis during fin regeneration. Whole mount in situ hybridization on a 3 dpa fin using a *dachc* probe to visualize distal niche cells (in blue). Blue arrowheads indicate peripheral fin regions with pronounced and broad *dachc* expression indicative of robust niche cell pools. Red arrowheads point to central fin tissue with modest *dachc* signal due to relatively small niche cell populations. The dashed red line marks the amputation plane.

**Supplemental Figure 9.**
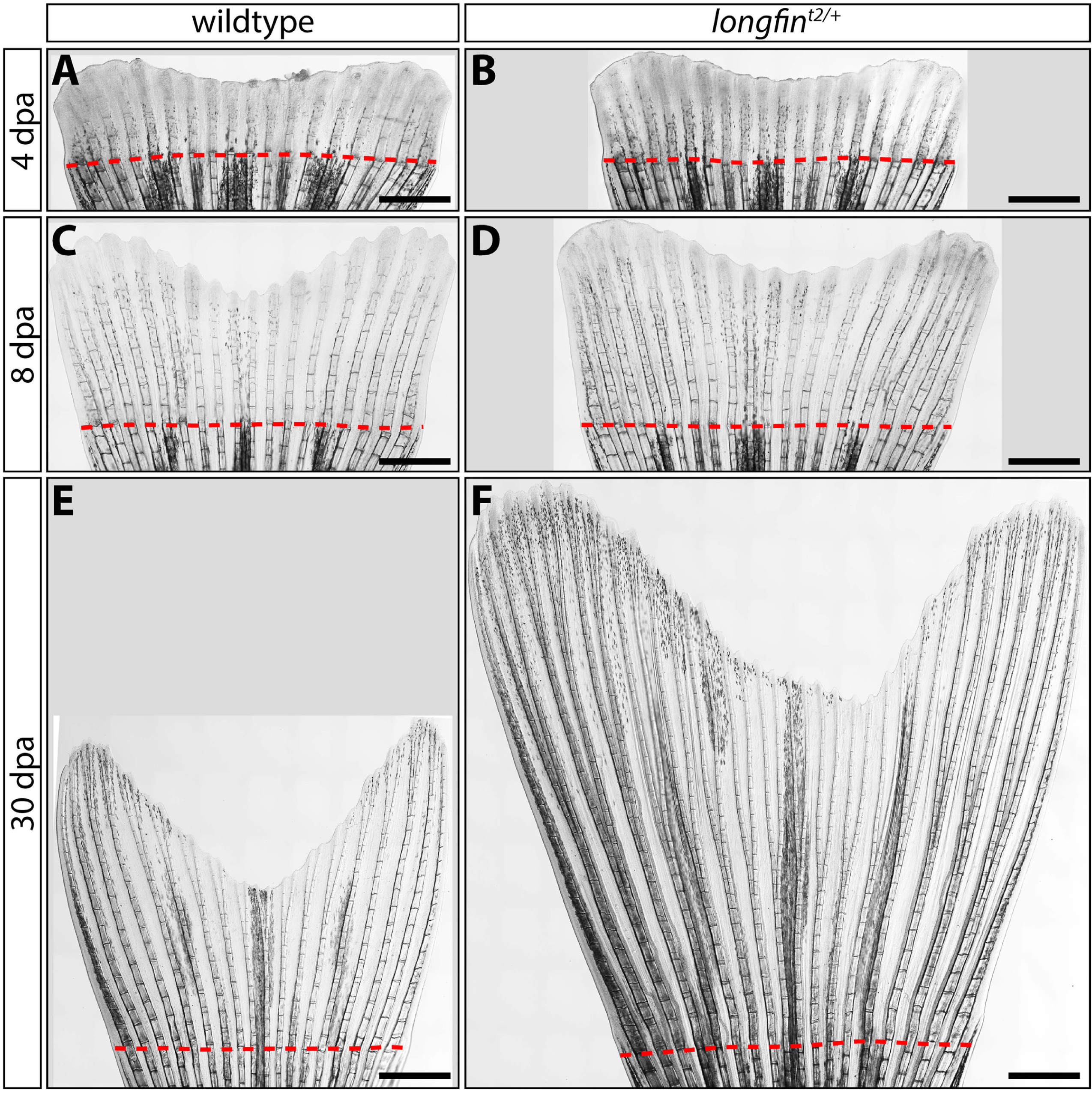
Regenerative outgrowth of *longfin^t2/+^*caudal fins fails to decelerate. **(A-F)** Stitched differential interference contrast (DIC) microscope images of caudal fins from clutchmate wildtype and *longfin^t2/+^* animals at the indicated day post amputation (dpa). Outgrowth is similar through 8 dpa but persists in *longfin^t2/+^*fish, leading to exceptionally long fins by 30 dpa. Quantitative data for the entire time course (n ≥ 12) is shown in Fig. 4. Dashed red lines highlight amputation planes. Scale bars represent 1 mm.

**Supplemental Figure 10.**
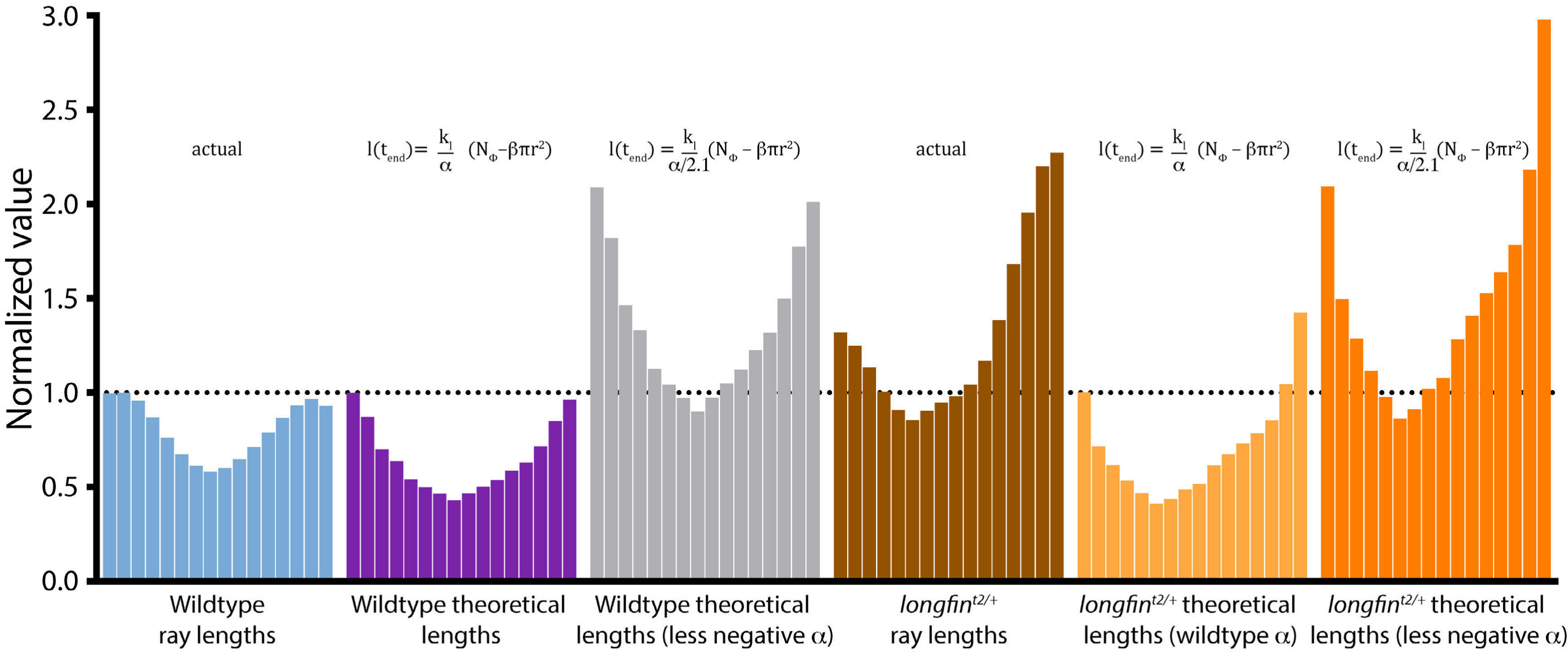
l*o*ngfin fin ray skeletal geometry and modeled regenerative outgrowth suggests *longfin* have deficient niche depletion – a broken “countdown timer”. Each distinctly colored set of distributed bars represents actual ray lengths or theoretical regenerated ray lengths for the 16 central rays (from a fixed proximal-distal position). All ray lengths are normalized to the longest wildtype ray. Theoretical lengths are derived from the transpositional scaling equation, which predicts final length as a function of ray radius, with scaling parameters set to restore the widest wildtype ray to its actual length. The alpha parameter (α) then is increased by 2.1x (by dividing α, a negative number, by 2.1; empirically set by linear regression comparing wildtype and *longfin* ray area to length relationships) to model a decreased rate of niche depletion that would cause wildtype resected rays to theoretically regenerate to a *longfin^t2/+^*length. Applying this same increase in α but inputting *longfin^t2/+^*ray radii into the transpositional scaling formula successfully predicts *longfin^t2/+^* fin-resected fish would regenerate with a *longfin* phenotype.

